# Sex differences in myocardial ketone oxidation capacity in cardiometabolic disease with coincident ischemic heart disease

**DOI:** 10.64898/2026.11.11.765904

**Authors:** Paramesha Bugga, Michael W. Stoner, Janet R. Manning, Bellina A.S. Mushala, Maryam Sharifi-Sanjani, Nisha Bhattarai, John Sembrat, Iain Scott

**Affiliations:** Vascular Medicine Institute, Department of Medicine, University of Pittsburgh, Pittsburgh, Pennsylvania, USA; Center for Metabolism and Mitochondrial Medicine, Department of Medicine, University of Pittsburgh, Pittsburgh, Pennsylvania, USA; Division of Cardiology, Department of Medicine, University of Pittsburgh, Pittsburgh, Pennsylvania, USA; Division of Pulmonary, Allergy, Critical Care, and Sleep Medicine, Department of Medicine, University of Pittsburgh, Pittsburgh, Pennsylvania, USA

**Keywords:** HFpEF, Myocardial Infarction, Ketone Oxidation, Metabolism, Cardiometabolic Disease

## Abstract

**Background:** Myocardial infarction (MI) in patients with heart failure with preserved ejection fraction (HFpEF) is associated with worse outcomes than MI in individuals without HFpEF. However, the molecular mechanisms underlying this increased vulnerability remain poorly understood. We investigated sex differences in cardiac ketone body (KB) metabolism in patients with diabetes-associated ischemic heart disease, and in a mouse model of HFpEF complicated by MI (HFpEF-MI).

**Methods:** Cardiac BDH1 and OXCT1 expression was assessed in male and female patients with diabetes, with or without ischemic heart disease. Male and female mice received a high-fat diet and L-NAME for 10 weeks to induce HFpEF, followed by coronary artery ligation to induce MI. Glucose tolerance, cardiac function, survival, cardiac KB oxidative capacity, and circulating KB levels were evaluated.

**Results:** Cardiac BDH1 and OXCT1 abundance was significantly reduced in male patients with diabetes-associated ischemic heart disease but preserved in female patients. In mice, 10 weeks of high-fat diet and L-NAME administration increased fasting blood glucose and impaired glucose tolerance. MI reduced systolic function in control and HFpEF mice of both sexes. Male HFpEF-MI mice exhibited higher mortality than female HFpEF-MI mice. Consistent with our human results, male HFpEF-MI mice showed reduced cardiac ketolytic enzyme expression alongside elevated circulating KB levels, whereas female mice maintained cardiac KB oxidative capacity.

**Conclusion:** Reduced cardiac ketolytic enzyme expression and impaired KB oxidative capacity are associated with greater mortality in male HFpEF-MI mice, whereas female mice exhibit preserved KB oxidative capacity and improved survival. Similar sex differences in ketolytic enzyme expression were observed in patients with diabetes-associated ischemic heart disease. These findings identify cardiac ketone metabolism as a potential therapeutic target warranting further investigation in HFpEF complicated by MI.

## Introduction

Heart failure with preserved ejection fraction (HFpEF) is a heterogeneous clinical syndrome characterized by signs and symptoms of heart failure despite a left ventricular ejection fraction ≥50%, accompanied by evidence of elevated cardiac filling pressures or congestion (1, 2). The development and progression of HFpEF are closely associated with aging, obesity, hypertension, type 2 diabetes, and chronic kidney disease (1–3). These conditions collectively impair myocardial relaxation, increase ventricular stiffness, and compromise vascular function and cardiovascular reserve (3). Thus, preserved ejection fraction does not imply normal cardiac performance, particularly during exercise, when impaired ventricular filling and limited cardiac reserve contribute to dyspnea and exercise intolerance (4). Ischemic heart disease is an important comorbidity that may contribute to the development and progression of HFpEF (5, 6). Myocardial ischemia can result from obstructive epicardial coronary artery disease, coronary microvascular dysfunction, or both. A prospective study of hospitalized patients with HFpEF identified a high prevalence of obstructive coronary artery disease and microvascular dysfunction, including previously unrecognized abnormalities (5, 7). Similarly, the PROMIS-HFpEF study demonstrated a high prevalence of impaired coronary flow reserve in patients without known revascularized macrovascular coronary artery disease. These findings underscore the need to distinguish HFpEF accompanied by obstructive coronary artery disease from HFpEF characterized predominantly by microvascular dysfunction.

Impaired myocardial energetics may provide a mechanistic link between HFpEF and ischemic heart disease. Cardiac relaxation requires ATP to support calcium reuptake into the sarcoplasmic reticulum and actomyosin cross-bridge detachment (8). Ischemia may further compromise oxidative ATP production by restricting oxygen availability, potentially exacerbating preexisting energetic deficits. Altered substrate utilization may also limit the myocardium’s ability to meet its energy demands (9). Human myocardial metabolomic studies have identified disturbances in pathways involving fatty acids, amino acids, and other oxidative substrates in HFpEF (9); however, metabolite concentrations alone do not establish metabolic flux. Susceptibility to cardiometabolic and ischemic heart disease differs between men and women, and varies with age and menopausal status (10–12). The mechanisms underlying the identified sex differences remain incompletely understood, highlighting the need for further investigation to identify potential sex-specific therapeutic targets.

Ketone bodies (KBs) have attracted interest as alternative myocardial energy substrates (13–16). Nevertheless, increased circulating KB availability does not necessarily translate into greater cardiac KB oxidation or ATP generation for cardiac contraction and relaxation, particularly when ketolytic capacity is impaired. The enzymes β-hydroxybutyrate dehydrogenase 1 (BDH1) and succinyl-CoA:3-oxoacid CoA transferase 1 (OXCT1) mediate key steps in cardiac ketone utilization. In recent years, both ketolytic enzymes have been investigated for their cardioprotective effects in various cardiovascular diseases (17–19). However, the roles of these enzymes in sex-specific responses to cardiometabolic disease complicated by ischemic heart disease remain poorly understood. For the first time, we demonstrate how this pathway is altered in HFpEF complicated by MI and that these changes differ between sexes, which may clarify the metabolic basis of susceptibility to ischemic injury. Accordingly, we examined sex differences in cardiac ketolytic enzyme expression in patients with diabetes-associated ischemic heart disease and investigated their relationship with cardiac KB oxidative capacity, dysfunction, and survival in experimental HFpEF-MI

## Materials and methods

### Human tissue samples

Fresh human cardiac tissue samples were collected from the left ventricles of organ donors deemed not suitable for transplant, under a protocol approved by the University of Pittsburgh Committee for Oversight of Research and Clinical Training (CORID). Tissue samples were flash-frozen and stored at −80°C until processing. Study authors only had access to the de-identified tissue samples and their associated clinical phenotyping data, and no personally discernible information was obtained. Due to the difficulty of obtaining HFpEF biopsy tissue, we used tissues from chronic diabetic/obese (DIA) patients as a surrogate for HFpEF, and tissues from patients with diabetes/obesity and ischemic heart disease (DIA-IHD) as a surrogate for HFpEF-MI. These groups are matched in our preclinical mouse model detailed below.

### Animal studies

All mice were bred and conserved in compliance with protocols approved by the University of Pittsburgh Institutional Animal Care and Use Committee. Male and female mice were used to study sex differences in cardiometabolic diseases (HFpEF) with and without ischemic heart disease.

### HFD/L-NAME-induced heart failure with preserved ejection fraction (HFpEF)

An HFpEF phenotype was induced in both male and female C57BL/6J background mice by administering a high-fat diet (HFD; Research Diets D12492; containing 60% calories from fat) and L-NAME (Sigma) at 0.5 g/L in drinking water. The development of HFpEF was authenticated by quantifying increased body weight, insulin resistance, cardiac diastolic dysfunction (as presented by perturbed echocardiographic parameters e.g. E/e’ ratio, E/A ratio), left ventricular hypertrophy, and cardiac interstitial fibrosis. All experimental mice were euthanized according to approved protocols. Hearts and blood samples were collected and snap-frozen for biochemical analysis (20).

### LAD ligation surgery

The LAD surgery procedure was adopted from Bugga et al. 2025 and Xing et al. 2021 (21, 22). Mice were sedated with 3% isoflurane in an induction chamber, then intubated. Dorsal surface hair was removed by applying Nair; thoracotomy was performed between the 3rd and 4th intercostal ribs, and the heart was exposed using homemade retractors. The left descending coronary artery (LAD) was visualized below (∼2 mm) the left atrium, and the LAD occluded permanently with 8-0 suture. Ischemia was confirmed by blanching of the myocardium. Intercostal muscles and ribs were sutured back with 6-0 silk sutures, and the skin closed with 6-0 silk sutures in an interrupted pattern. Cardiac function and myocardial strain were measured after 7 days of MI surgery. For sham-operated groups, we followed a similar procedure except for coronary artery ligation.

### Intraperitoneal glucose tolerance test

Intraperitoneal glucose tolerance test (IPGTT) was performed on overnight-fasted mice. D-glucose was dissolved in sterile saline, and a glucose bolus (2 g/kg) was injected intraperitoneally. Blood glucose levels were measured before glucose solution injection (i.e. 0 min) and after 15, 30, 60, and 120 min. The data were analyzed, and an area under the curve was calculated (23).

### Echocardiography

Cardiac function was evaluated by transthoracic echocardiography using a Vevo 3100 system equipped with an MS400 transducer (Visual Sonics Inc). Anesthesia was induced by 3% isoflurane and confirmed by lack of response to firm pressure on one of the hind feet. Throughout echocardiogram acquisition, body temperature was controlled using a heating pad, and isoflurane was lowered to 1.0–1.5% and adjusted to preserve heart rate in the range of 350-450 bpm. Left ventricular ejection fraction and other indicators of systolic function were recorded from short-axis M-mode scans at the papillary muscle level under isoflurane-induced anesthesia in gently restrained mice. Apical 4-chamber views were used to evaluate diastolic function measurements using pulse-wave and tissue Doppler at the level of the mitral valve. The parasternal long-axis view was used to quantify global longitudinal strain (GLS), and speckle tracking analysis was employed to confirm GLS of the myocardium. Parameters gathered include: heart rate (HR), left ventricular end-diastolic diameter (LVIDd), left ventricular end-systolic diameter (LVID,s), end-diastolic interventricular septal wall thickness (IVSd), left ventricular end-diastolic posterior wall (LVPWd), LV mass, left ventricular fractional shortening (LVFS), left ventricular ejection fraction (LVEF), peak Doppler blood inflow velocity across the mitral valve during early diastole (E), peak Doppler blood inflow velocity across the mitral valve during late diastole (A), isovolumic relaxation time (IVRT), and peak tissue Doppler of myocardial relaxation velocity at the mitral valve annulus during early diastole (e’). At the end of the procedures, all mice recovered from anesthesia without difficulties. All parameters were measured at least 3 times, and averages are presented.

### Serum ketone body measurement

Blood samples were collected from all the experimental animals under anesthesia by the cardiac puncture method. Serum was collected by centrifuging the samples at a speed of 2500 rpm for 10 min. Samples were frozen at -80^0^ C until use. Serum ketone bodies were measured using a commercially available kit (Cat No. 700190; Cayman Chemicals). Briefly, a 1:2 dilution of each serum sample was prepared using the BDH assay buffer, and 50 μL of test and samples were loaded into a 96-well plate. Next, 50 μL of detection reagent was added to the wells, and the samples were incubated at RT for 30 min. The absorbance was measured at 450 nm using a plate reader, and the concentration of BHB was calculated by extrapolating the OD values to the standard curve.

### Immunoblotting

Protein samples were prepared from frozen mouse heart tissue by lysis in ice-cold CHAPS lysis buffer (150 mM NaCl, 50 mM Tris HCL pH 7.4, 1% Triton X-100, 0.5% sodium deoxycholate, 0.1% SDS, 5 mM EDTA) containing protease and phosphatase inhibitors. Proteins were separated by SDS-PAGE on 4-12% or 12% gradient gels (Invitrogen) and transferred to nitrocellulose membranes. An Odyssey scanner (LI-COR version 3.0) was used as an exposure system. Protein expression was quantified using a fluorescently labelled secondary antibody, and protein band intensities were analyzed with ImageJ software. Proteins were detected with the following primary antibodies from Protein Tech: GAPDH, Cat No: 60004-1-1g; 1:5000; BDH1, Cat No: 15417-1-AP; 1:2000; OXCT1, Cat No: 12715-1-AP; 1:5000.

### Statistical analysis

All data are represented as means ± SEM. Statistical analyses of studies with two variables were shown using two-way ANOVA followed by Tukey’s post hoc test. Studies involving a single variable were evaluated with unpaired, two-tailed Student’s t tests. A *P* value ≤ 0.05 was considered significant, and all analyses were performed in GraphPad Prism 10.0. All experiments were performed with at least three biological replicates. Experimental animals were randomly allocated to each group. Investigators were blinded to the various experimental conditions of the individual animals during both the experiments and the experimental analysis. Comparison of survival rates was performed by Kaplan-Meier analysis.

## Results

### Cardiac ketone oxidative capacity compromised among cardiometabolic disease-associated ischemic heart disease patients

Cardiac KB oxidation plays an important role in cardiometabolic health (17, 19), and circulating KB levels are usually high among the diabetes population (14). We measured cardiac KB oxidation enzyme abundance in control, diabetes, and diabetes-associated ischemic heart disease patients (**Figure 1A**). Beta-hydroxybutyrate dehydrogenase (BDH1) and Succinyl-CoA:3-ketoacid coenzyme A transferase 1 (SCOT1/OXCT1) were significantly decreased among male diabetes-associated ischemic heart disease patients but preserved in female patients from the same group (**Figure 1B**). This data suggests that cardiac ketone oxidative capacity is significantly compromised among male patients with diabetes-associated ischemic heart disease, and may promote worse outcomes after an ischemic event.

**Figure 1:**
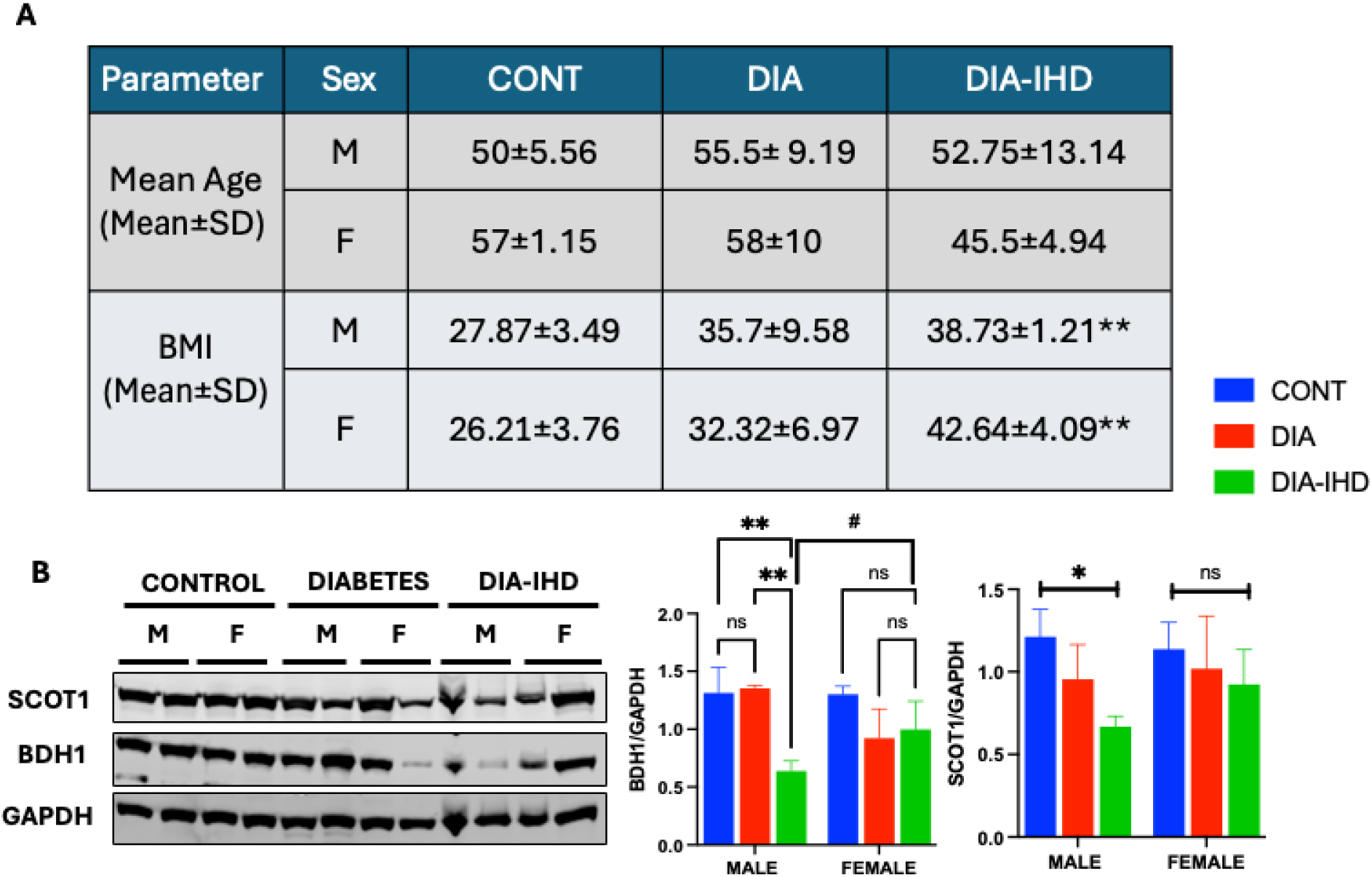
Ketone body oxidative capacity is reduced among male patients with diabetes-associated ischemic heart disease. **A.** The table shows age and BMI of control, diabetes, diabetes ischemic heart disease patients. B. Western blot image of BDH1 and OXCT1/SCOT1 proteins. N = 3. Data are represented as Mean ± SEM. *p<0.05 vs. Control, **P<0.01 vs. Control, ^#^P<0.05 Male vs. Female in DIA-IHD. CONT; Control, DIA; Diabetes, DIA-IHD; Diabetes-associated ischemic heart disease.

### HFD/L-NAME feeding for 10 weeks successfully increased glucose intolerance and fasting blood glucose levels

To induce the HFpEF phenotype, 10-to 12-week-old male and female mice were fed a high-fat diet (HFD; 60% of total calories from fat) and received L-NAME (0.5 g/L) in drinking water for 10 weeks. Body weight was monitored throughout the treatment period to assess changes associated with this regimen. Both male and female mice exhibited significant weight gain during treatment (**Figure S1A,B**). Following 10 weeks of treatment, we assessed glucose homeostasis by measuring fasting blood glucose and performing an intraperitoneal glucose tolerance test. HFD/L-NAME treatment significantly increased fasting blood glucose levels in both male and female mice compared with their respective controls, indicating impaired fasting glucose regulation (**Figure 2C,F**). Male mice exhibited higher fasting blood glucose levels than female mice, suggesting a more pronounced disturbance in glucose homeostasis. To further characterize glucose handling, mice received an intraperitoneal glucose injection (2 g/kg), and blood glucose concentrations were measured at baseline (0 min) and 15, 30, 60, and 120 min after injection. The resulting data were used to calculate the area under the curve as an integrated measure of the glycemic response. Male mice receiving HFD/L-NAME exhibited greater glucose intolerance than similarly treated female mice, as reflected by a higher glucose area under the curve (**Figure 1A-F**). Together, these findings demonstrate that HFD/L-NAME treatment was associated with weight gain and impaired glucose homeostasis in both sexes, with male mice exhibiting more pronounced fasting hyperglycemia and glucose intolerance.

**Figure 2:**
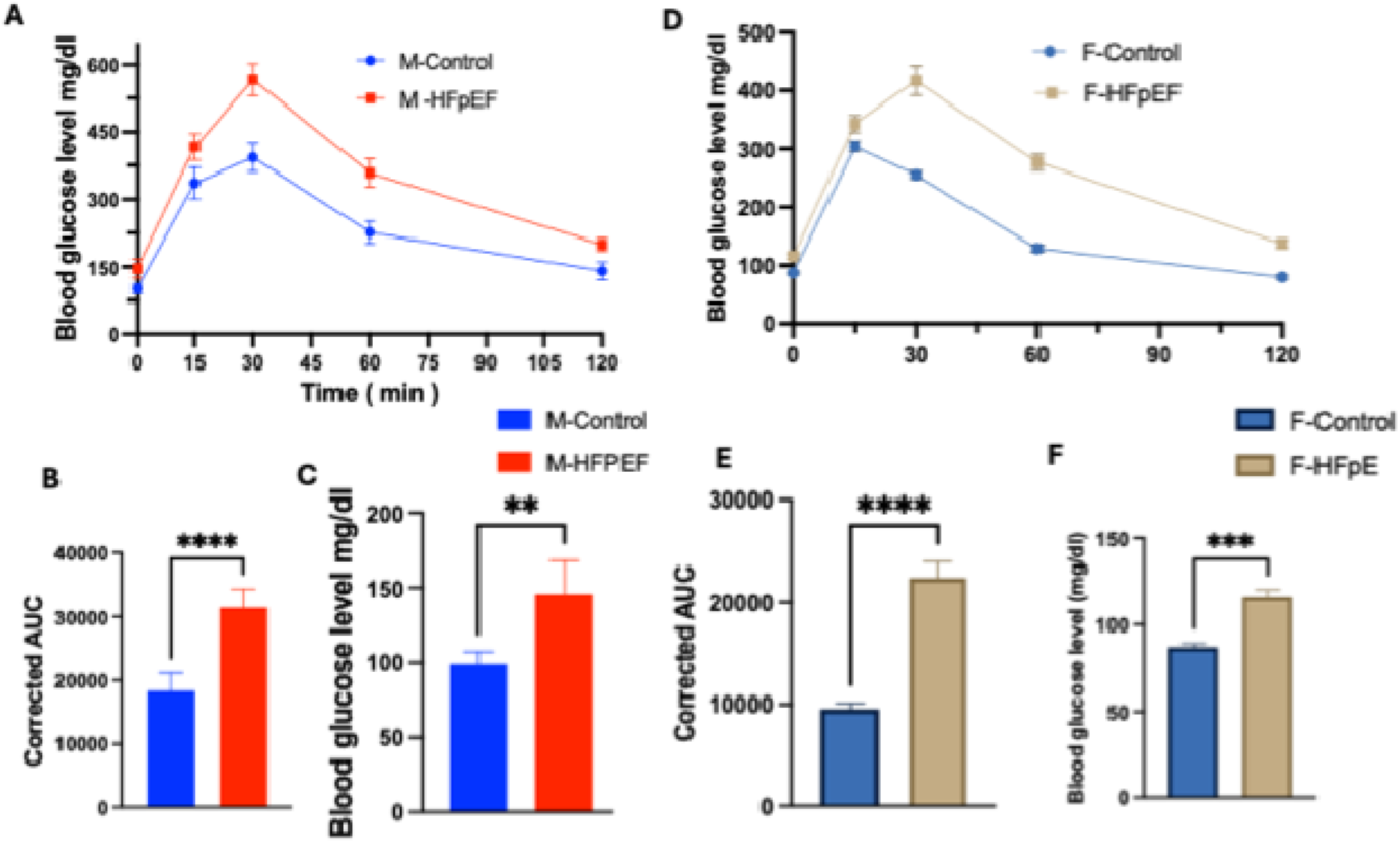
HFD/LNAME administration successfully induced glucose intolerance in male and female mice. **A.** Intraperitoneal glucose tolerance test (IPGTT) of male animals. **B.** Area under the curve of blood glucose levels of males. **C**. Fasting blood glucose levels of males. **D.** IPGTT of female animals. **E**. Area under the curve of blood glucose levels of females. **F**. Fasting blood glucose levels of females. N = 5. Data are represented as Mean ± SEM. **P<0.01 vs. Control, ***P<0.001 vs. Control, ****P<0.0001 vs. Control, M: Male, F: Female, HFpEF: heart failure with preserved ejection fraction.

### HFD/L-NAME feeding for 10 weeks resulted in diastolic dysfunction in both male and female mice

Next, we assessed cardiac systolic and diastolic function using echocardiography. We found that both male and female mice fed with HFD/L-NAME had preserved cardiac systolic function (**Figure 3A-B, E-F; Figure S2B-C,E-F**), and observed a significant increase in LV mass in both male and female mice (**Figure S2A,D**). We also assessed diastolic function by measuring mitral valve flow and mitral annular velocity using pulse wave and tissue Doppler methods, respectively. Both male and female animals showed diastolic dysfunction after HFD/L-NAME feeding, as shown by increased E/e’ ratios (**Figure 3C,G**).

**Figure 3:**
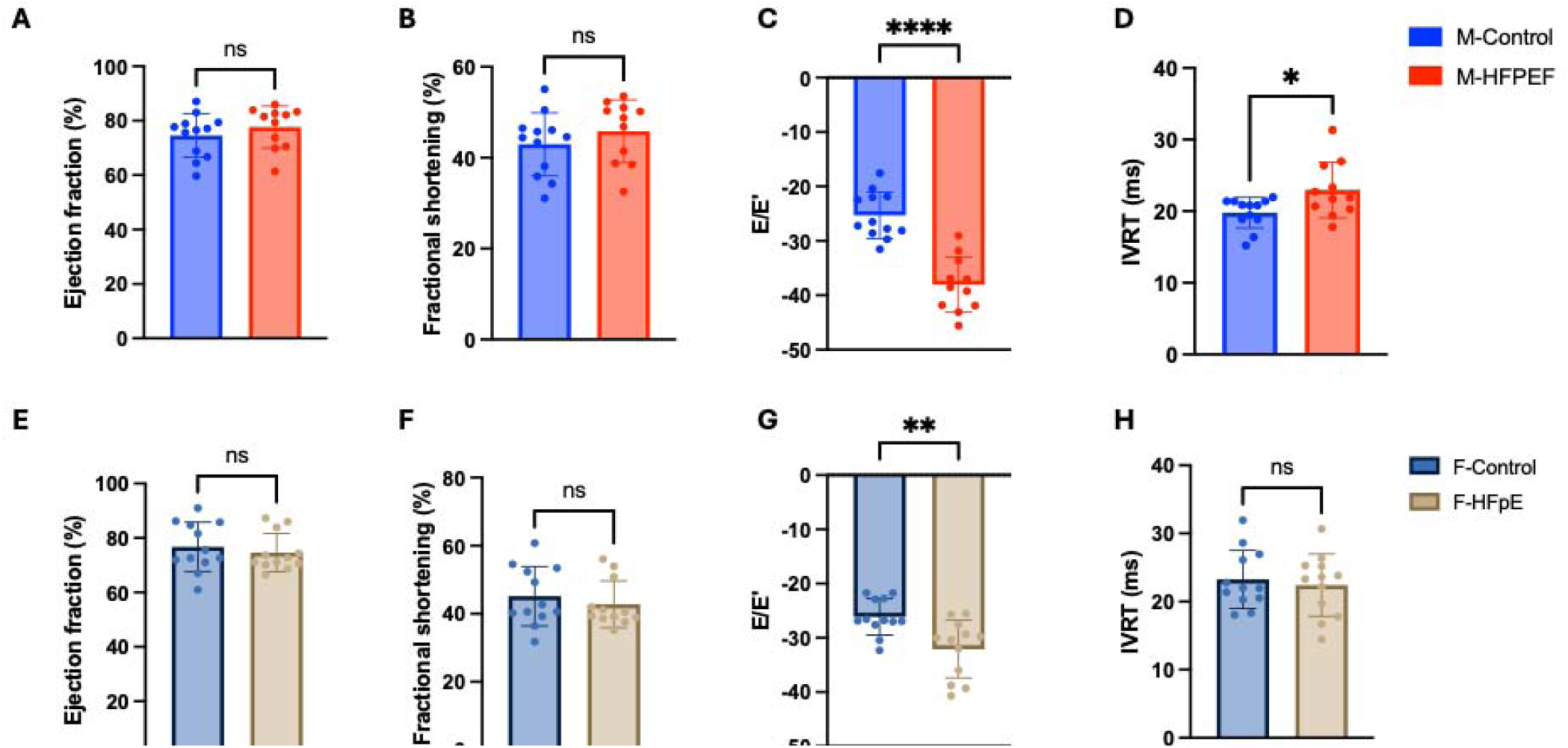
HFD/L-NAME feeding successfully induced cardiac diastolic dysfunction in male and female animals without affecting systolic function. A. Ejection fraction of male mice. B. Fractional shortening of male mice. C. E/e’ ratio in male mice. D. Isovolumic relaxation time in male mice. E. Ejection fraction of female mice. F. Fractional shortening of female mice. G. E/e’ ratio in female mice. H. Isovolumic relaxation time in female mice. Data are represented as Mean ± SEM. *P<0.05 vs. Control, **P<0.01 vs. Control, ***P<0.001 vs. Control, M: Male, F: Female, HFpEF: heart failure with preserved ejection fraction.

Although both male and female animals exhibited diastolic dysfunction, we witnessed a more prominent increase in E/e’ and IVRT parameters (*P* < 0.001 vs. control, and *P* < 0.01 vs. control, respectively) in males than in females (*P* < 0.01 vs. control, and n.s. vs. control, respectively) after HFD/L-NAME feeding, similar to the increase in LV mass. Together, we conclude that while cardiac dysfunction was induced in both male and female mice, this finding was more severe in male animals.

### Myocardial infarction followed by HFpEF induction leads to significantly higher mortality in male mice

After establishing cardiac diastolic dysfunction, we performed LAD ligation surgery (i.e., MI surgery) in male and female HFpEF mice. Seven days after surgery, we evaluated cardiac function by echocardiography in control-SHAM, control-MI, HFpEF-SHAM, and HFpEF-MI male and female mice. We noted a significant reduction in cardiac systolic function after MI surgery in both male and female mice (**Figure 4A-C**). We also observed a significant preservation of cardiac systolic function in control-MI female mice in comparison to control-MI male animals (**Figure 4A-C**).

**Figure 4:**
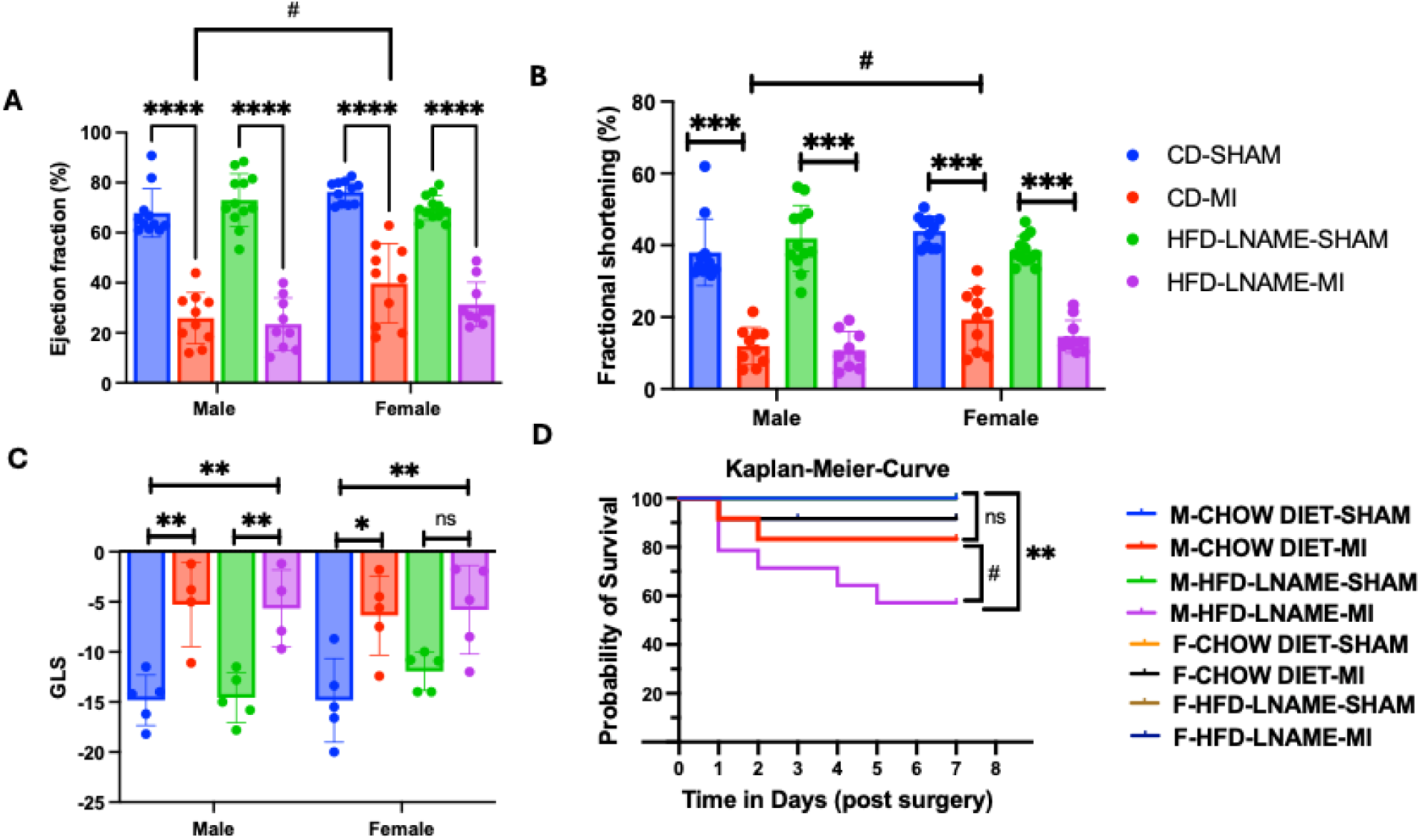
HFpEF-associated myocardial infarction leads to a significantly higher mortality in male mice. **A.** Ejection fraction of male and female mice post-MI surgery**. B.** Fractional shortening of male and female mice post-MI surgery. **C.** Global longitudinal strain analysis of male and female mice post-MI surgery. **D.** Survival curve analysis of male and female mice after surgery. Data are represented as Mean ± SEM. *P<0.01 vs. SHAM, **P<0.01 vs. SHAM , ***P<0.001 vs. SHAM, ****P<0.0001 vs. SHAM, # p<0.05 vs. Male MI vs. Female MI, M: Male, F: Female, HFpEF: heart failure with preserved ejection fraction.

Survival analysis was performed after MI surgery in all experimental animals using Kaplan–Meier curve analysis. We observed significantly higher mortality in the male HFpEF-MI group compared with the male control-SHAM, with a total of 6 mortality events (**Figure 4D**), whereas the control-MI group had 3 deaths (**Figure 4D**). Only one death was observed in the female HFpEF-MI group animals, demonstrating that female mice were more protected than males under the same condition (**Figure 4D**).

### Cardiac ketone body oxidative capacity is reduced in HFpEF-MI male animals and preserved in female mice

Finally, we wanted to identify the potential pathological mechanism responsible for the significant mortality observed with HFpEF-associated myocardial infarction in male mice. Because we observed a significant decrease in cardiac ketone oxidative enzyme expression (i.e., BDH1 and SCOT1) in male diabetes-associated ischemic heart disease, we assessed cardiac ketone body oxidative enzyme expression in male and female mice after HFpEF and HFpEF-MI stress. We noticed a significant decrease in cardiac KB oxidative enzyme expression BDH1 and SCOT1 levels in male MI only (*P* < 0.05), HFpEF (*P* < 0.05), and HFpEF-associated MI (*P* < 0.01), compared with the control-SHAM group of animals (**Figure 5A-C**). In contrast, ketone oxidation enzyme expression levels were preserved among female mice under the same experimental conditions as male mice (**Figure 5A-C**).

**Figure 5:**
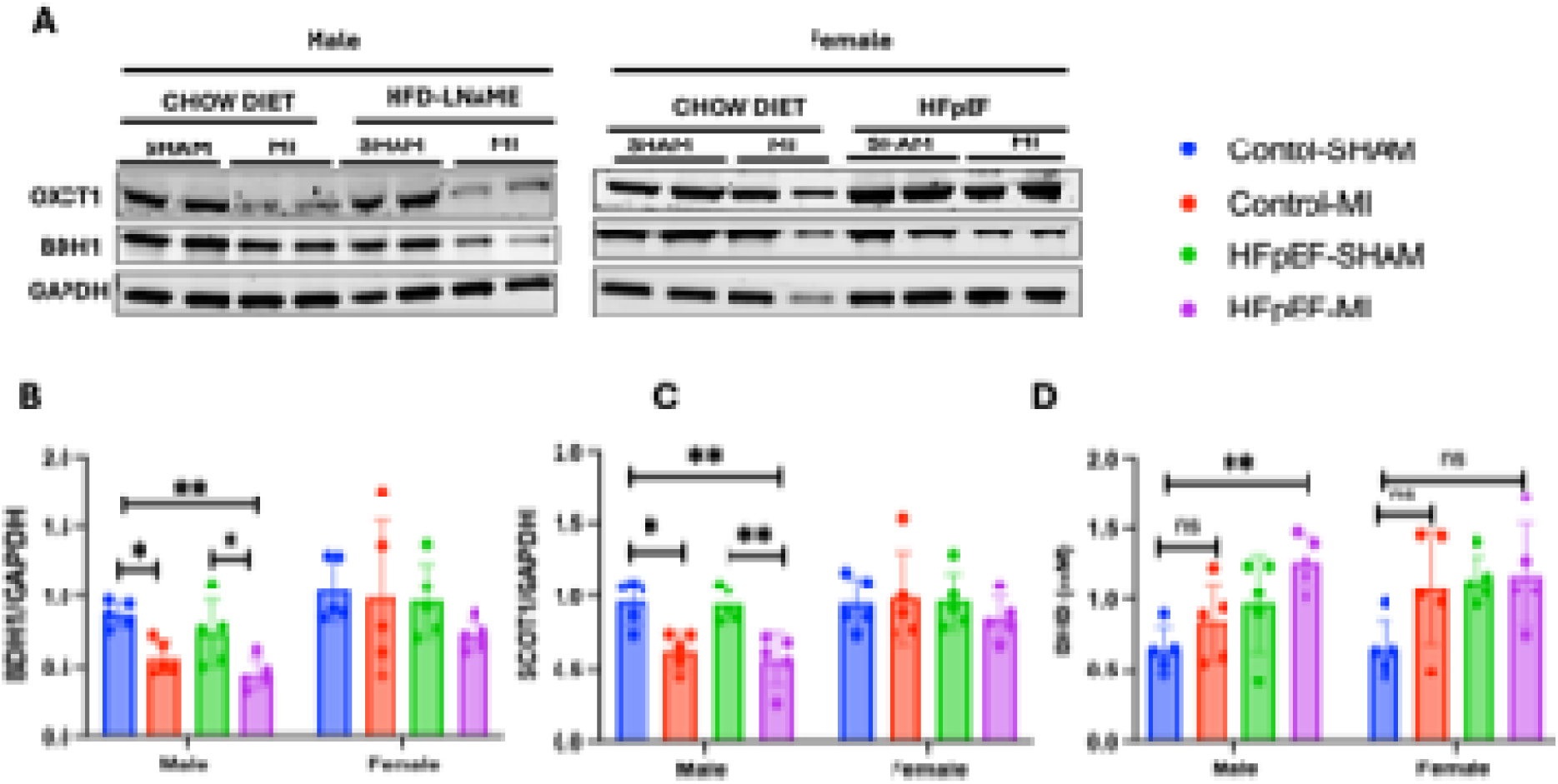
Female mice display preserved cardiac KB oxidation capacity and lower ketone body accumulation than male mice. **A.** Representative images of cardiac BDH1 and SCOT1/OXCT1 protein expression in male and female mice. **B.** Cardiac BDH1 expression in male and female mice. **C.** Cardiac SCOT1/OXCT1 expression in male and female mice. **D.** Circulating BHB levels in male and female mice 1-week post-surgery N = 5. Data are represented as Mean ± SEM. *p<0.05, **p< 0.01.

Next, we quantified the circulating levels of KB in all the experimental animals post-surgery, to determine the potential effect of decreased cardiac KB oxidation enzyme. In line with our earlier findings, circulating KB were increased in animals with decreased cardiac KB oxidative enzyme expression, and male animals with HFpEF-MI showed a significant increase in circulating KB levels compared to control-SHAM male animals (**Figure 5D**). Female animals from MI, HFpEF and HFpEF-MI groups showed moderate, but non-significant increases in serum KB levels in comparison to control female animals (**Figure 5D**).

## Discussion

Coronary artery disease is more prevalent among the HFpEF patient population compared to non-HFpEF controls (25), and both cardiometabolic disease and ischemic heart disease affect men and women differently (11, 26, 27). Circulating KB levels are high among HFpEF, diabetes, and diabetes-associated ischemic heart disease patients (28–30). However, despite having high circulating KB levels, patients with diabetes and HFpEF are vulnerable to cardiovascular complications. We therefore hypothesized that compromised cardiac ketolytic capacity may contribute to the high vulnerability to cardiac complications among HFpEF and HFpEF-associated ischemic heart disease patients. For the first time, we demonstrate the potential role of cardiac ketone oxidation capacity in the pathophysiology of HFpEF and HFpEF-associated ischemic heart disease in both male and female patients and experimental animals. We found a significant decrease in cardiac ketone body-oxidizing enzymes (i.e., BDH1 and OXCT1) in male patients with diabetes-associated ischemic heart disease. Consistent with our findings, Deng et al. (2021) demonstrated that decreased cardiac BDH1 expression and increased KB levels were responsible for cardiac inflammation in a 3-hit HFpEF model (16). In a recent report, Wang et al. (2026) revealed the cardioprotective activity of BDH1; cardiac-specific BDH1 overexpression mitigated mitochondrial damage by activating the AKT/GSK-3β pathway in a STZ- and high fat diet-induced diabetic cardiomyopathy model (31). Shi et al. (2026) showed that cardiac-specific BDH1 overexpression protects mice from streptozotocin-induced diabetic cardiomyopathy by inhibiting cardiomyocyte apoptosis (32). Finally, reports by Brahma et al. and Cia et al. drew similar conclusions that the cardiac KB oxidizing enzymes BDH1 and OXCT1 were significantly lower in chronic diabetes and heart failure patients, and in STZ- and highfat diet-induced preclinical diabetes animal models (33, 34).

Consistent with Tong et al. (35), female mice in our study exhibited less glucose intolerance and milder diastolic dysfunction than male mice, suggesting lower susceptibility to cardiometabolic HFpEF, and further supporting sex-dependent differences in disease vulnerability. In contrast to our findings, Cao et al. reported that female mice are more susceptible to cardiometabolic disease than male mice (36). This discrepancy could be due to mouse strain differences or the age difference of the experimental mice at the start time point. Our novel preclinical HFpEF and HFpEF-associated myocardial ischemia revealed an interesting finding: we observed greater mortality among male mice (i.e., 50%) in the HFpEF-MI group than in the female HFpEF-MI group (i.e., 25%), without a significant difference in cardiac function. Although cardiac systolic function was not statistically significantly higher in HFpEF-MI females (i.e., EF; 28.6%) than in HFpEF-MI males (i.e., EF; 22.6%), female mice showed preserved cardiac systolic function, and this minor preservation might be responsible for lower mortality in the female HFpEF-MI group. On the other hand, control-MI male mice presented statistically significant systolic dysfunction compared with control-MI female mice. This could be due to healthy females showing greater resistance to MI-induced myocardial damage than the HFpEF female mice.

The lower mortality observed in HFpEF-MI female mice compared with HFpEF-MI male mice inspired us to investigate whether sex differences in myocardial ketone metabolism might contribute to these outcomes. Consistent with our observations in human myocardial samples and findings from others, cardiac BDH1 and OXCT1/SCOT protein expression were significantly reduced in male mice subjected to MI, HFpEF, or HFpEF-MI, whereas female mice maintained ketolytic enzyme expression across these conditions. These findings suggest that female mice retain the enzymatic machinery required for myocardial ketone utilization during cardiometabolic and ischemic stress. Female control-MI mice showed significantly better-preserved systolic function and no observed deaths compared with male control-MI mice. This difference may be related to preserved cardiac ketolytic capacity in females, which was significantly reduced in males after MI. Aubert et al. reported increased cardiac BDH1 in a heart failure mouse model, transverse aortic constriction (TAC)-MI (37). However, the experimental models varied: we induced ischemia following metabolic stress, whereas their study induced ischemia after pressure overload caused by TAC. The studies also used mice of different ages, which may have contributed to the differences in outcomes. We found that circulating KB levels increased in males one week after surgery and showed an upward trend in male mice across the MI, HFpEF, and HFpEF-MI groups, with a significant increase in HFpEF-MI males relative to male controls, consistent with previous clinical reports of diabetes and heart failure events (30, 38). Female mice exhibited modest increases in circulating KB levels in MI, HFpEF, and HFpEF-MI experimental conditions. The coexistence of elevated circulating ketones and reduced cardiac ketolytic enzyme expression in males is consistent with a potential mismatch between ketone availability and myocardial utilization. Together, these findings support sex-dependent differences in cardiac ketone oxidative capacity that may contribute to exacerbation of cardiac damage and lead to mortality due to HFpEF-associated MI stress among the male mice.

## Conclusion

Collectively, our findings demonstrate for the first time that loss of cardiac ketolytic enzyme expression occurs in male mice with MI, HFpEF, or HFpEF-MI and male patients with diabetes-associated ischemic heart disease. In contrast, female mice and female patients with diabetes-associated ischemic heart disease maintained cardiac ketone oxidative capacity. Female mice also exhibited lower mortality than males after MI and HFpEF-MI cardiac stress. Restoring the cardiac KB oxidative capacity can be a potential therapeutic target to reduce mortality associated with HFpEF-associated ischemic heart disease in men and postmenopausal women.

## Limitations of the study

We considered diabetes and diabetes associated ischemic disease human cardiac tissue samples as HFpEF and HFpEF-associated ischemic heart disease conditions due to the limited availability of human HFpEF samples. HFpEF is an asymptomatic disease, and obtaining human HFpEF samples is challenging. Diabetes and obesity are the key comorbid conditions; we considered the diabetic patients’ samples as HFpEF samples. We demonstrated decreased cardiac KB oxidative capacity by downregulation of BDH1 and OXCT1 protein expression. Further, validation with more clinical samples is needed to confirm this by quantifying the cardiac KB levels and BDH1 and OXCT1 enzymatic activity.

## Author Contributions

P.B. and I.S. conceived the study and designed experiments. P.B. executed experiments. P.B. analyzed data. P.B. and I.S. prepared figures. M.W.S., J.R.M., B.A.S.M., N.B., and M.S.-S. provided critical input and expertise. P.B. and I.S. drafted the manuscript. P.B. and I.S. edited and revised the manuscript. All authors approved the final submission.

## Acknowledgments

This work was supported by National Institute of Health Fellowships (F31DK134089 and T32HL110849) to B.A.S.M. and National Institute of Health Research Grants (R01HL147861, R0HL156874) and an American Heart Association Established Investigator Award (23EIA1037834) to I.S. Echocardiography was carried out by the University of Pittsburgh Rodent Ultrasonography Core, which received funding from the NIH Shared Instrumentation Grant Program (S10OD023684).

## Supplementary Data

## Supplementary Data

**Fig. S1.**
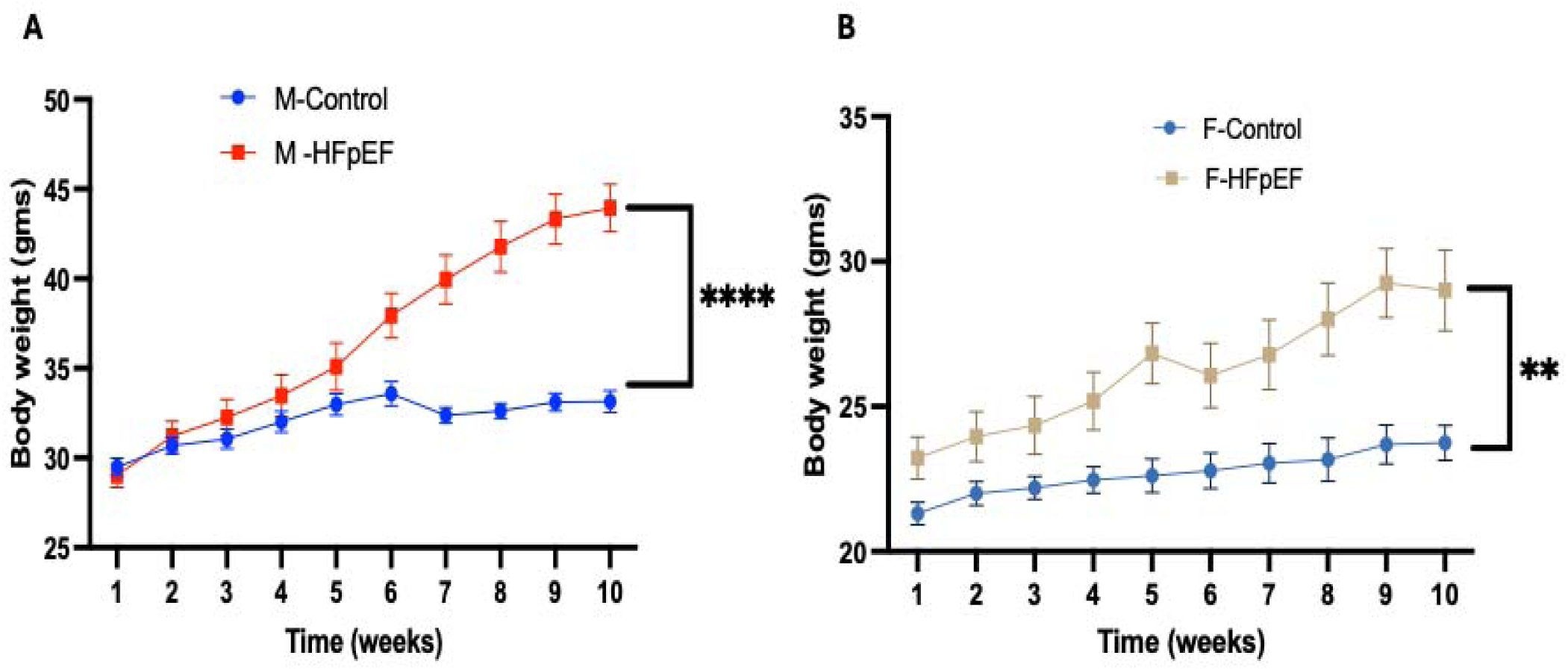
**HFD-LNAME administration significantly increases body weight of both male and female animals**. *A. Overall body weight change of male animals during the study duration. B. Overall body weight change of female animals during the study duration. Data were represented as Mean±SEM. **P<0.01 Vs Control, ***P<0.001 Vs Control, ****P<0.0001 Vs Control, M: Male, F: Female, HFpEF: Heart Failure with preserved Ejection Fraction*.

**Fig. S2.**
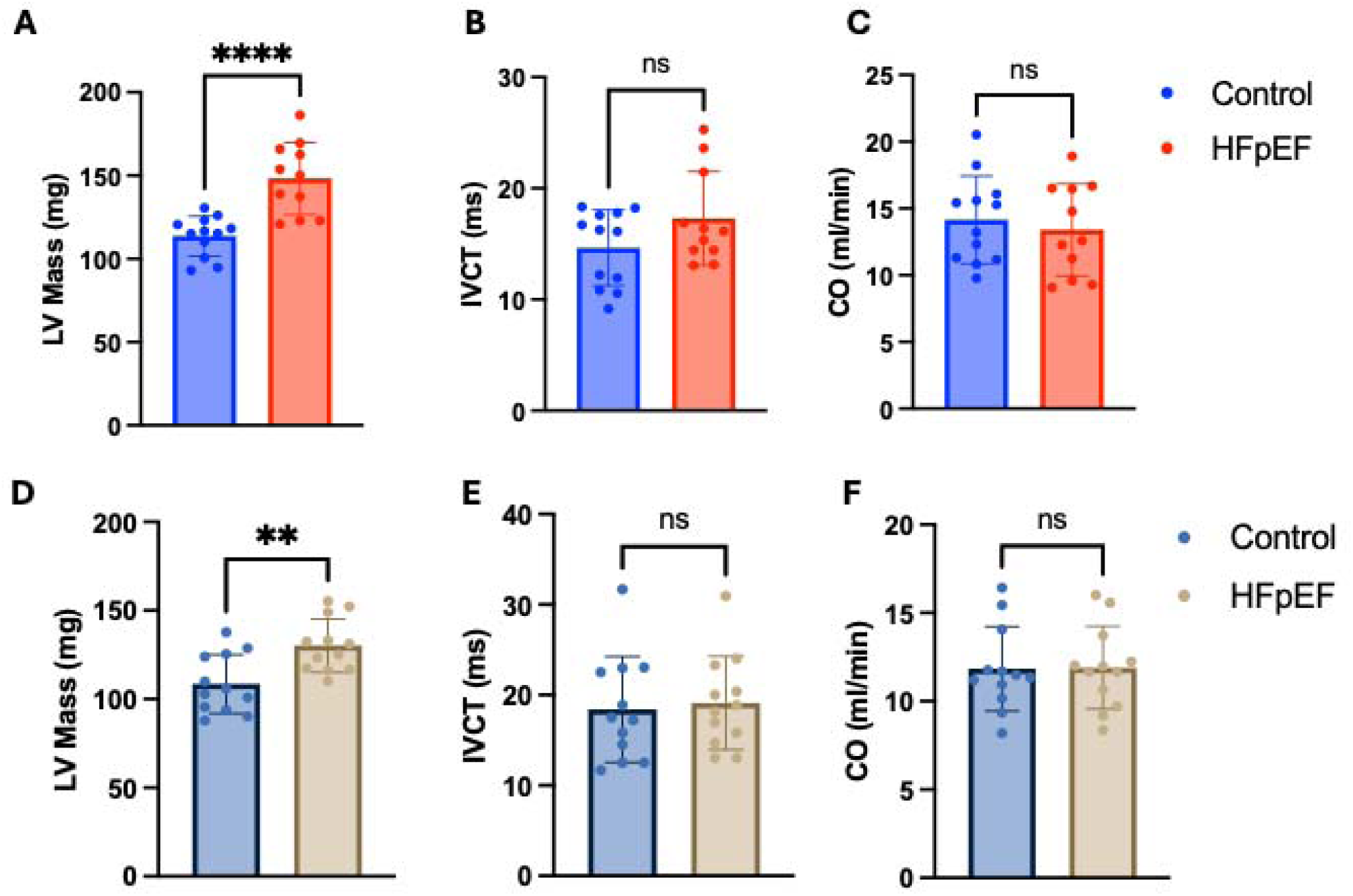
**HFD-LNAME feeding successfully induced diastolic dysfunction in male and female animals without affecting the systolic function of the heart**. A. Left ventricular Mass of male mice. *B. Isovolumic contraction time of male mice. C. Cardiac output of male mice. D. Left ventricular Mass of female mice. E. Isovolumic contraction time of male mice. F. Cardiac output of female mice. Data were represented as Mean±SEM. **P<0.01 Vs Control, ***P<0.001 Vs Control, M: Male, F: Female, HFpEF: Heart Failure with preserved Ejection Fraction*.

